# Morphologic characterization and cytokine response of chicken bone-marrow derived dendritic cells to infection with high and low pathogenic avian influenza virus

**DOI:** 10.1101/2024.02.06.579192

**Authors:** Jongsuk Mo, Karen Segovia, Klaudia Chrzastek, Kelsey Briggs, Darrell R. Kapczynski

**Affiliations:** Exotic and Emerging Avian Disease Research Unit, U.S National Poultry Research Center, Agricultural Research Service, USDA, 934 College Station Road, Athens, GA 30605, USA; CSL Seqirus, 255 Wyman Street, Waltham, MA 02139, USA; Animal and Plant Health Agency, Pathology and Animal Sciences, APHA, Addlestone KT15 3NB, UK

**Keywords:** Chicken, innate immunity, dendritic cells, avian influenza, cytokines, interferon

## Abstract

Dendritic cells (DCs) are professional antigen-presenting cells, which are key components of the immune system and involved in the early immune response. DCs are specialized in capturing, processing, and presenting antigens to facilitate immune interactions. Chickens infected with avian influenza virus (AIV) demonstrate a wide range of clinical symptoms, based on pathogenicity of the virus. Low pathogenic avian influenza (LPAI) viruses typically induce mild clinical signs, whereas high pathogenic avian influenza (HPAI) induce more severe disease, which can lead to death within days. For this study, chicken bone marrow-derived DC (ckBM-DC)s were produced and infected with high and low pathogenic avian influenza viruses of H5N2 or H7N3 subtypes to characterize innate immune responses, study effect on cell morphology, and evaluate virus replication. A strong proinflammatory response, including chicken interleukin-1β, and stimulation of the interferon response pathway were observed at 8 hours post infection. Microscopically, the DCs underwent morphological changes from classic elongated dendrites to a more general rounded shape that eventually lead to cell death with the presence of scattered cellular debris. Differences in onset of morphologic changes were observed between H5 and H7 subtypes. Increases in viral titers demonstrated that both HPAI and LPAI are capable of infecting and replicating in DCs. The elevated expression of infected DCs may be indicative with a dysregulation of the immune response typically seen with HPAI infections.

## 1. Introduction

In recent years, avian influenza virus (AIV) has been one the leading causes of infection-based poultry mortality and morbidity. Prior to the 1990s, AIV outbreaks in domesticated poultry were rare, however ongoing outbreaks of highly pathogenic avian influenza (HPAI) have occurred globally for the past several years [1–3]. The H5 A/goose/Guangdong/1996 (H5-Gs/Gd) lineage is responsible for most of the outbreaks, as the current clade 2.3.4.4b viruses appear to be highly adapted to migratory waterfowl [3]. As a result of the adaptation, more spill over into domesticated poultry, mammals, and humans have been observed [4]. High morbidity and mortality rates have led to reduced poultry production, embargoes on countries of origin, and increased expenses associated with vaccinating and controlling AIV within the global poultry industry [5, 6]. In 2022, a total of 67 countries reported HPAI outbreaks, resulting in the deaths of 131 million poultry and wild birds [7, 8]. In the U.S., the ongoing 2022-2023 HPAI H5N1 outbreak has resulted in the loss of over 60 million birds and $3 billion dollars in economic damages [1, 9].

Low pathogenic avian influenza (LPAI) viruses typically cause a mild disease in poultry that is restricted to the respiratory and intestinal tract because they contain a mono-basic cleavage site in the hemagglutinin (HA) protein that can only be cleaved by a few, localized cellular proteases [6, 10, 11]. HPAI viruses contain a multi-basic cleavage site that allows for several common proteases to cleave the HA, which leads to a severe, systemic infection. [6, 11]. The rapid, multi-organ infection coupled with a HPAI-specific dysregulated cytokine response typically leads to death 1-6 days post infection in domesticated poultry [12]. Early responses against viral infections are pre-dominantly mediated by host innate immunity. Increased expression of pathogen recognition receptors (PRRs), interferons, pro-inflammatory cytokines, and chemokines are generally observed during the early stages of an AIV infection [13]. PRRs, such as Toll-like receptors (TLRs) and MDA-5, sense viral RNAs and initiate an inflammatory response by releasing proinflammatory cytokines. A rapid induction of type 1 (interferon-alpha (IFN-α)) and type 2 (interferon-beta (IFN-β)) interferon leads to the upregulation of interferon stimulated genes (ISG)s, which are essential for an antiviral response. One ISG, myxovirus resistance gene (Mx) is important because it promotes anti-AIV activity in various mammalian and avian species [14–16]. Proinflammatory cytokines, including interleukin 6 (IL-6), interleukin 12 (IL-12), and interleukin 1 beta (IL-1β) are released upregulate the inflammatory response to help fight the infection. AIV can also stimulate cell death or apoptosis, which acts as a host-defense mechanism to inhibit viral replication in host cells, and is mediated by the enzymes, Caspase-3 (Casp-3) and Caspase-8 (Casp-8) [17–23]. The expression of these innate immune modulators drastically varies by virus strain, host, and target tissue making our understanding of the immune response to AIV incomplete [12, 13].

Antigen presenting cells (APC) are crucial components of the primary immune response against pathogens and help the bridge the innate and adaptive immune responses. Dendritic cells (DC) are professional APCs, that play a central role as regulators of the adaptive immune response by interacting with T and B cells [13]. In the case of mammals, DCs and macrophages are usually within proximity of each other, which enables them to form a network capable of rapidly countering foreign antigens and react against inflammatory stimuli [24]. While different in terms of structure and morphology, avian species are also known to have a similar network of DCs and macrophages that react swiftly against foreign antigens [25]. DC progenitors originate from hematopoietic stem cells in the bone marrow and translocate to non-lymphoid tissue where they become immature DCs [26]. While immature DCs are capable phagocytizing antigens, they are poor T-cell stimulators lacking proper antigen presentation capabilities. Upon activation, they migrate to T-cell regions where they mature and express several costimulatory molecules, including MHC-II, CD11c, CD40 and CD80. Mature DCs are specialized in antigen presentation to T cells [27]. Recently, more emphasis has been put on understanding the immune modulation of chicken DC cells and their ability to combat disease. The use of prebiotics has been used to increase the activity of DCs and therefore, T and B cells, and studies have examined the expression profile of various cytokines originating from chicken DCs infected with other poultry diseases such as Newcastle disease virus [28, 29].

One early study reported DCs can be grown by co-culturing chicken bone marrow (BM)-derived cells with chicken granulocyte-macrophage stimulating factor (GM-CSF) and chicken interleukin 4 (IL-4) [30]. In this study, we culture bone-derived chicken dendritic cells (ck-BM-DC) to examine gene expression levels of IFN-α, TLR-3, TLR-7, MHC-I, IL-1β, IL-6, Mx, Casp-3, and Casp-8 of DCs infected with AIV There are limited studies examining the interactions between chicken DCs and AIV [25, 31–33]. The exact nature of how AIV infections affect DCs is largely unknown. We seek to determine whether active AIV replication can occur in DCs and if antigen processing occurs. In this study, we compared immune responses, morphological changes, and replication of ckBM-DCs following infection with contemporary H5 and H7 HPAI and LPAI viruses. A better understanding of how chicken antigen presentation occurs is needed, as the Gs/Gd lineage becomes entrenched in migratory waterfowl globally.

## 2. Methods

### 2.1 Chickens and chicken bone marrow dendritic cells (ckBM-DC) isolation and culture

Four-week-old specific pathogen-free (SPF) white leghorn chickens were housed at the USDA-ARS-Southeast Poultry Research Laboratory (SEPRL). All birds used in these studies were cared for and handled in compliance with the Institutional Animal Care and Use Committee (IACUC) guidelines and procedures. ckBM-DCs were generated as previously described with minor modifications to the protocol [34]. Briefly, following euthanasia, femurs of chickens were removed and placed into 10 cm petri dishes containing PBS with 1% antibiotics (Sigma-Aldrich, St. Louis, MO). Both ends of the femur bone were cut across the tops with sterile bone-scissors and a sterile iron wire was passed through and the bone marrow was flushed with sterile PBS using 20 ml syringe with 16G needle. Marrow clusters were gently meshed through a 70 nm screen using a syringe plunger to obtain single-cell suspensions. Cell suspensions were overlaid with an equal volume of Histopaque 1119 (Sigma-Aldrich, St. Louis, MO) and centrifuged at 1200 g for 30 min at RT to remove red blood cells. Cells were collected and were washed three times in RPMI-1640 media (Thermo-fisher Scientific, Waltham, MA), before cells were counted by trypan blue exclusion assay.

Cells were cultured in six-well plates at a concentration of 2×10^6^ cells/ml at 41°C and 5% CO_2_ in RPMI-1640 supplemented with 10% chicken serum (Thermo-fisher, Waltham, MA), 1% L-glutamine, 1% non-essential amino acids and antibiotics (Gibco, Thermo-fisher, Waltham, MA) for 7 days. Different concentrations (0, 10, 25 and 50 ng/ml) of commercial IL-4 and commercial GM-CSF (Kingfisher, St Paul, MIM) were added to the medium to optimize culture conditions. Three-quarters of the medium was replaced with pre-warmed fresh complete medium at every 2 days. Cells were stimulated with LPS (500 ng/ml) (Thermo-fisher Scientific, Waltham, MA) for 30 hours to induce maturation. Images of the cells were taken at 30 hours using an EVOS 5000 (Invitrogen, Carlsbad, CA).

### 2.2 Virus propagation and inactivation

A total of 7 viruses, HPAI H5N2 A/chicken/Pennsylvania/1370/1983, H7N3 A/chicken/Jalisco/CPA1/2012 and LPAI H5N2 A/chicken/ Pennsylvania/21525/1983, A/Cinnamon Teal/Mexico/2817/2006 (H7N3), A/turkey/Virginia/SEP-4/2009 (H1N1) and A/Turkey/Wisconsin/68 (H5N9) strains were propagated in the allantoic cavities of 9–11 day old SPF chicken eggs. Viral titers were determined as previously described [35]. The H5N9 strain was inactivated with 0.01% β-propiolactone (BPL) overnight, followed by dialysis with sterile PBS. Inactivation of the virus was tested by performing serial passages on eggs. After inactivation, the strain was labeled with FITC labeling kit according to manufacturer’s recommendation (Thermo-fisher, Waltham, MA.)

### 2.3 Morphology and phenotypic analysis

Cells were cultured for 6 days in the presence of different concentrations (0, 10, 25 and 50 ng/ml) of GM-CSF and IL-4. Cell morphology and cell growth were monitored daily. After stimulation with LPS (500ng/ml) for 30 hours, images were taken from cell cultures to check change in morphology. Immunofluorescence labelling was performed to analyze the DC markers using FITC labeled mouse-anti-chicken MHC-II in combination with mouse anti-chicken CD40 (Bio-Rad, Hercules, CA), mouse anti-chicken CD11c (8F2, IgG2a) followed by incubation with a goat anti-mouse-Ig secondary Ab (Thermo-fisher Scientific, Waltham, MA). Primers for surface markers (MHC-II, CD40, CD11c, CD80, CD83 and CD86) were designed according to methods we have previously used [21].

### 2.4 Phagocytosis assay

Phagocytosis was assessed using FITC labeled inactivated H5N9 virus, and 0.5 um carboxylate modified fluorescent red latex beads (Sigma-Aldrich, St. Louis, MO). To explain briefly, non-stimulated ckBM-DCs were cultured for 6 days, followed by incubation with FITC-labeled inactivated H5N9 virus or chicken serum-opsonized red latex beads in RPMI-1640 medium at a density of 10^8^ particles/ml at 41°C for 4 hours. Cells were washed five times with PBS and visualized with immunofluorescence microscopy.

### 2.5 Immunofluorescence analysis

For sialic acid receptor staining, cells were fixed and stained by incubating FITC-labeled MAA (SA-α2,3-Gal) and TRITC-labeled SNA (SA-α2,6-Gal) for 1 hour at room temperature. Following 3 rinses in PBS, cells were stained for 5 minutes in DAPI (all reagents were purchased from Thermo-fisher Scientific, Waltham, MA). The immunofluorescence assays for virus nuclear protein (NP) detection were performed as previously described [36]. Briefly, cells were infected with A/turkey/Virginia/SEP-4/2009 (H1N1) and A/Turkey/Wisconsin/68 (H5N9) virus at a MOI of 1 for 20 hours. Cells were then washed with PBS twice, fixed and permeabilized with methanol. Viral antigens were detected with mouse-derived monoclonal antibody specific for a type A influenza virus nucleoprotein (developed at Southeast Poultry Research Laboratory, USDA), then stained with FITC-conjugated anti-mouse IgG antibody (Thermo-fisher Scientific, Waltham, MA).

### 2.6 Virus infection and analysis of cytokine expression by quantitative real-time RT-PCR (qRT-PCR)

Cells were infected with either LPAI or HPAI H5N2 and H7N3 at a MOI of 1 in serum free DC medium for one hour with gentle agitation applied every 10 minutes. Cells were washed twice with PBS and resuspended in DC medium containing 2% chick serum and incubated at 41°C and 5% CO2. At 2, 8 and 24 hours post infection (hpi), supernatants were collected and stored at -80°C until titration. Virus titers are expressed as log10 50% embryo infectious dose (EID_50_/ml) and HAU. Cells were harvested for RNA extraction at 8 hpi. Relative gene expression levels of IFN-α, Mx, TLR-3, TLR-7, MHC-I, IL-1β, IL-6, Casp-3 and Casp-8 were evaluated by qRT-PCR as previously described [21].

### 2.7 Statistical analyses

Data are expressed as the mean ± standard error. Statistical differences were analyzed with Tukey one-way ANOVA using Prism 9 (GraphPad Co., San Diego, CA).

## 3. Results

### 3.1 Morphological characteristics of chicken bone marrow-derived DC (ckBM-DC)

Morphological characteristics of ckBM-DC differed based on the levels of recombinant chicken GM-CSF and IL-4 (0, 10, 25 and 50 ng/ml) used. While no international consensus exists on how to determine units of activity for avian cytokines, we used 50 ng/ml of GM-CSF and 50 ng/ml IL-4 to attempt to maximize the number of cell aggregates, which were best observed at day 6 (Fig. 1. D). There was little difference among cells treated with 10 ng/ml or 25 ng/ml (Fig. 1. B, C). Non-treated ckBM-DCs demonstrated a more prominent rounded morphology, compared to cells treated with GM-CSF and IL-4, with the cells eventually detaching from the plate, leaving only a few live cells by day 6 (Fig. 1. A).

**Fig. 1.**
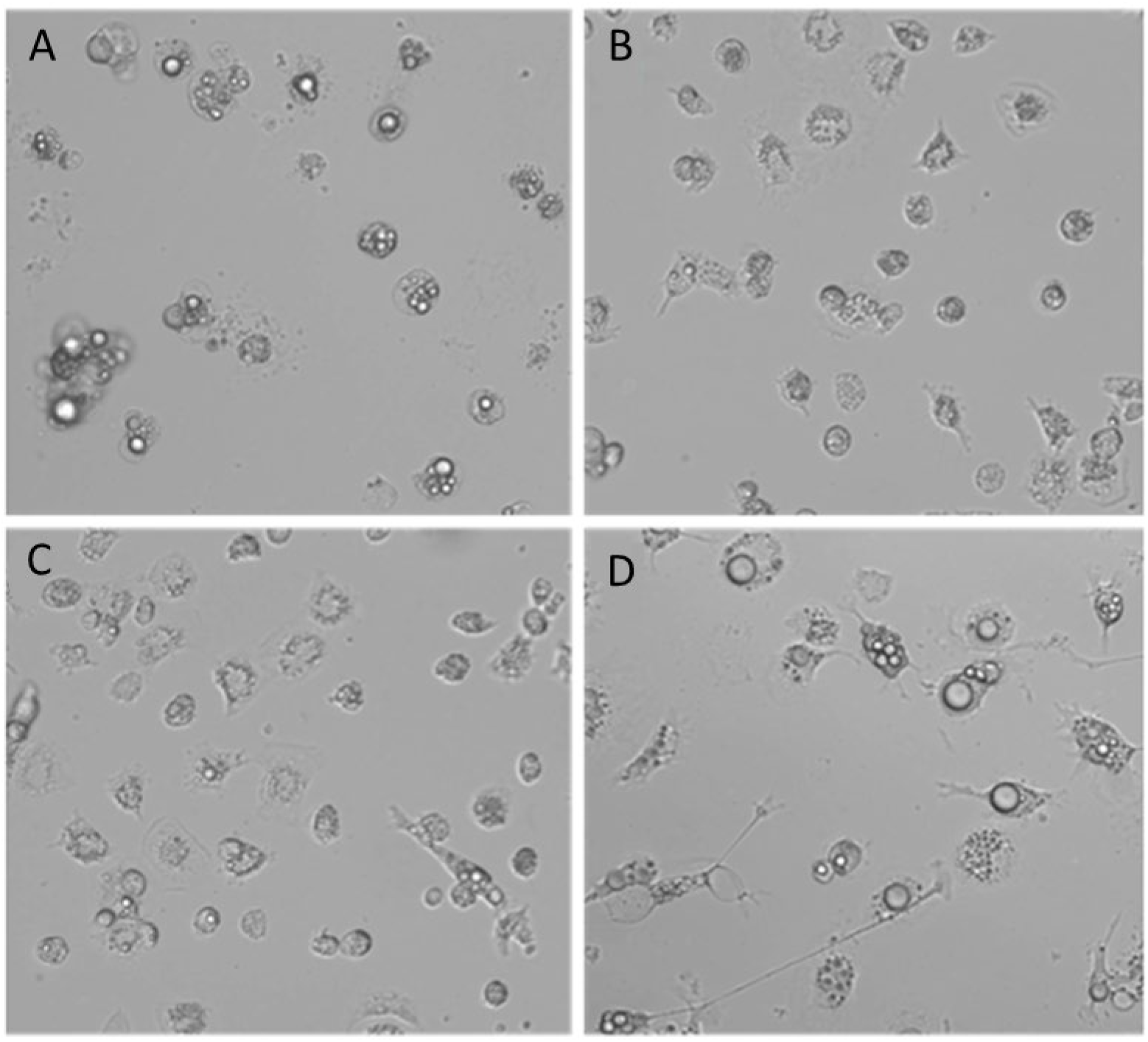
Morphology of ckBM-DC. Bone marrow-derived cells were cultured in the presence of different levels of recombinant chicken granulocyte-macrophage stimulating factor (GM-CSF) and recombinant chicken interleukin (IL)-4 for 6 days and dendrite formation was observed by microscopy. (A) 0 ng/ml GM-CSF+0 ng/ml IL-4. (B) 10 ng/ml GM-CSF+10 ng/ml IL-4. (C) 25 ng/ml GM-CSF+25 ng/ml IL-4. (D) 50 ng/ml GM-CSF+50 ng/ml IL-4. A representative image is shown for each concentration at 200x magnification.

### 3.2 Maturation of ckBM-DC

The ckBM-DCs were cultured in the presence of 50 ng/ml GM-CSF and 50 ng/ml IL-4. To induce maturation, we stimulated cells with 500 ng/ml LPS on day 6 for 30 hours. The cells were examined at different time points (0, 10, 20 and 30 hours). At 0 time point, cells displayed a veiled appearance, in which case they possessed large cytoplasmic veils (Fig. 2. A). When the cell aggregates were incubated with LPS for 10 hours, cell mophology changed with cells displaying long and thin branch-like features, with a spiny or sheet-like appearance. (Fig. 2. B). At the later timepoints,20 and 30 hours, more and more cells developed a dendritic-like appearance, indicating maturation of the cells. (Fig. 2. C, D).

**Fig. 2.**
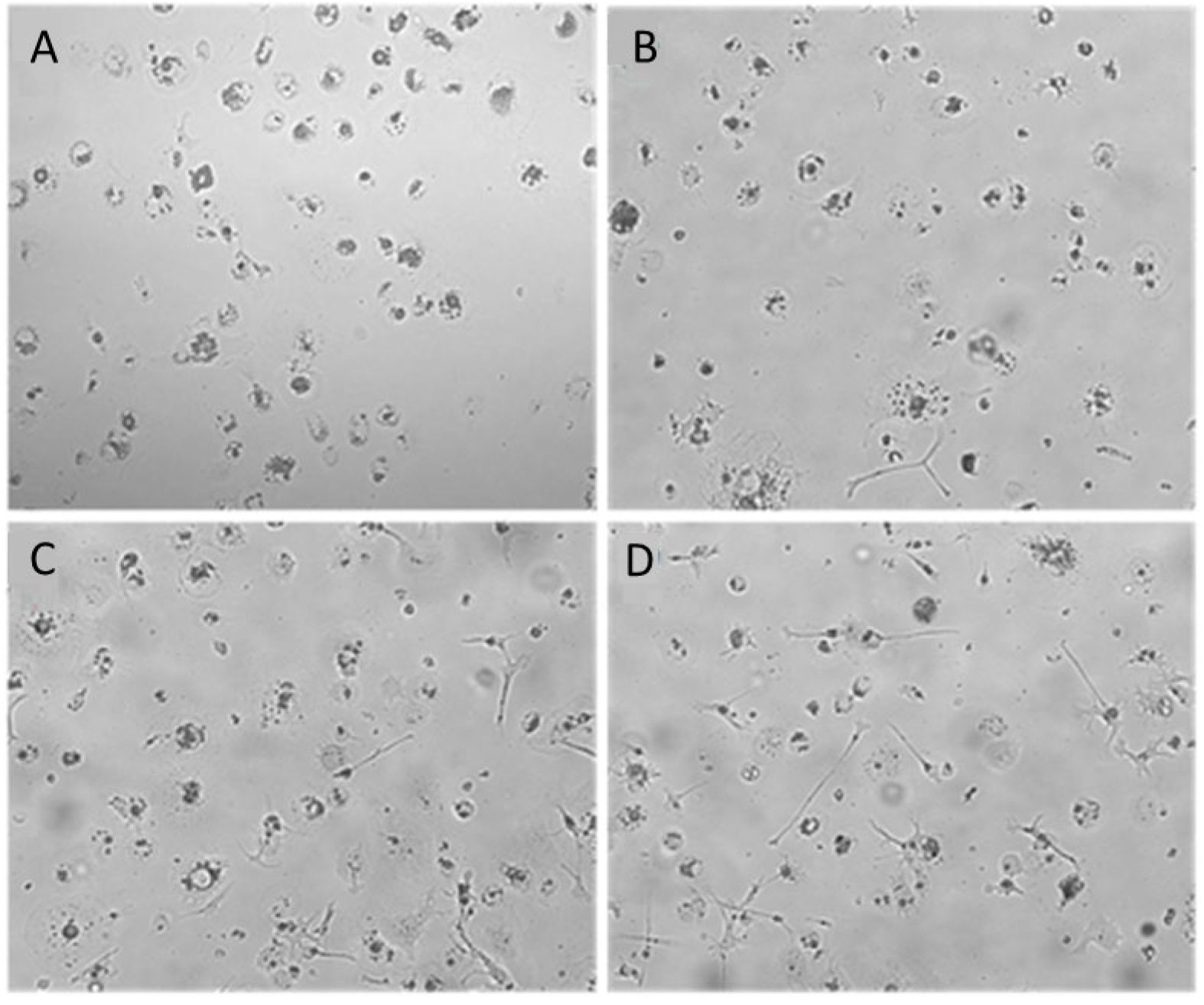
Morphology of ckBM-DC stimulated with LPS. Cells were cultured in the presence of 50 ng/ml GM-CSF + 50 ng/ml IL-4 for 6 days and then stimulated with LPS (500ng/ml). ckBM-DCs were observed by microscopy for 30 hours. Images show cells cultured at (A) 0 hour. (B) 10 hours. (C) 20 hours. (D) 30 hours. A representative image is shown for each timepoint at 100x magnification.

### 3.3 Mature ckBM-DC cells share phenotypic similarities with mammalian DC cells

Dendritic cells co-exist in both immature and mature states. In mammals, immature dendritic cells are characterized by moderate or low-level expression of surface markers molecules such as MHC-II, CD11c, CD40, CD80, CD83 and CD86 [24]. After stimulation with LPS, expression levels of all surface markers were increased. Immunofluorescence microscopy demonstrated that immature ckBM-DCs had some level of surface marker expression when stained with anti-chicken MHC-II (Fig. 3. A1), anti-chicken CD11c (Fig. 3. B1) and anti-chicken CD40 (Fig. 3. C1). After stimulation with LPS for 24 hours, expression level was increased in all 3 markers, MHC-II (Fig. 3. A2), CD11c (Fig. 3. B2) and CD40 (Fig. 3. C2). The level of surface marker expression was significantly enhanced in mature ckBM-DC cells, approximately 40-120-fold compared with their immature counterparts, especially CD80, CD83 and CD86 (Fig. 3. B)

**Fig. 3.**
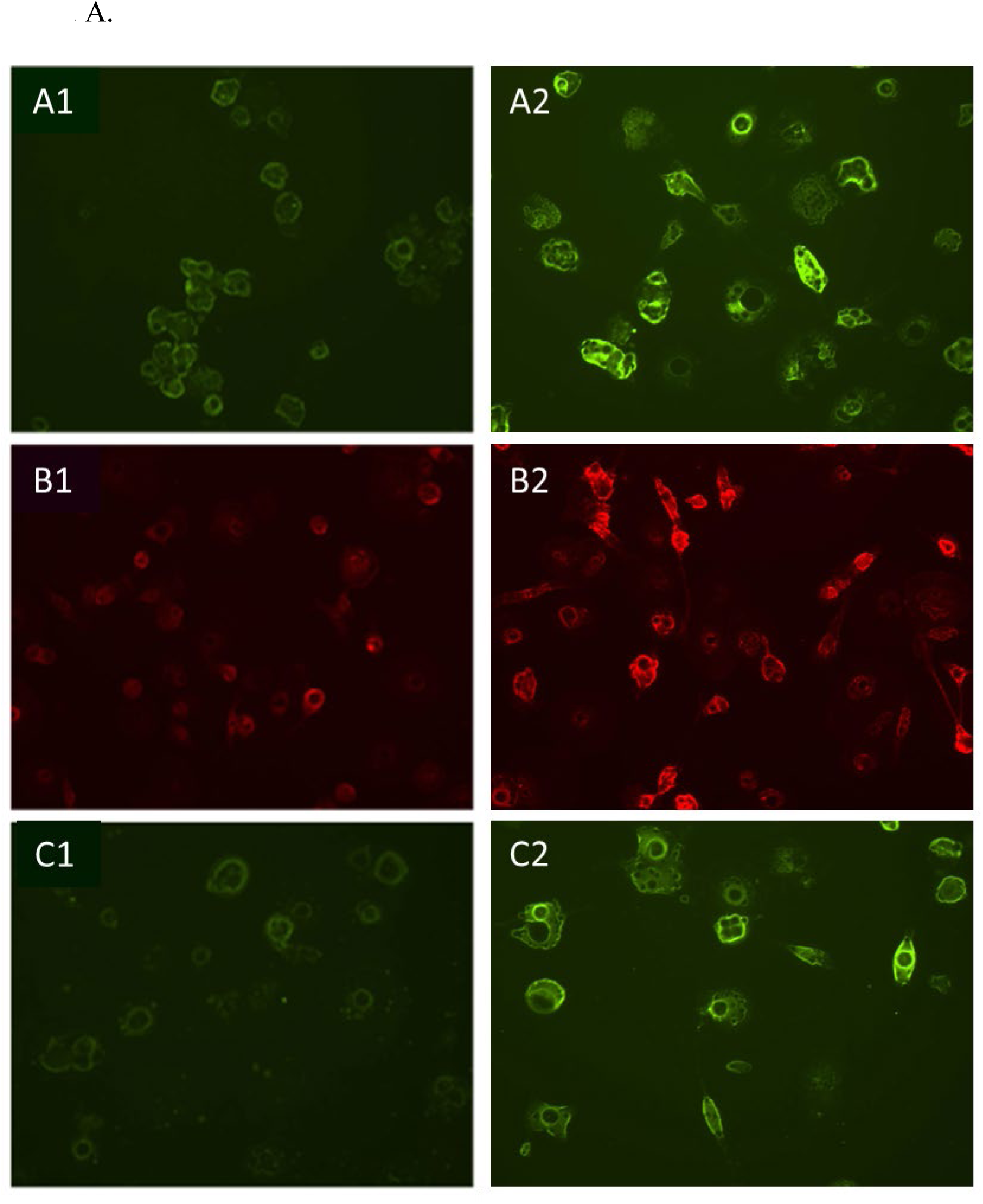
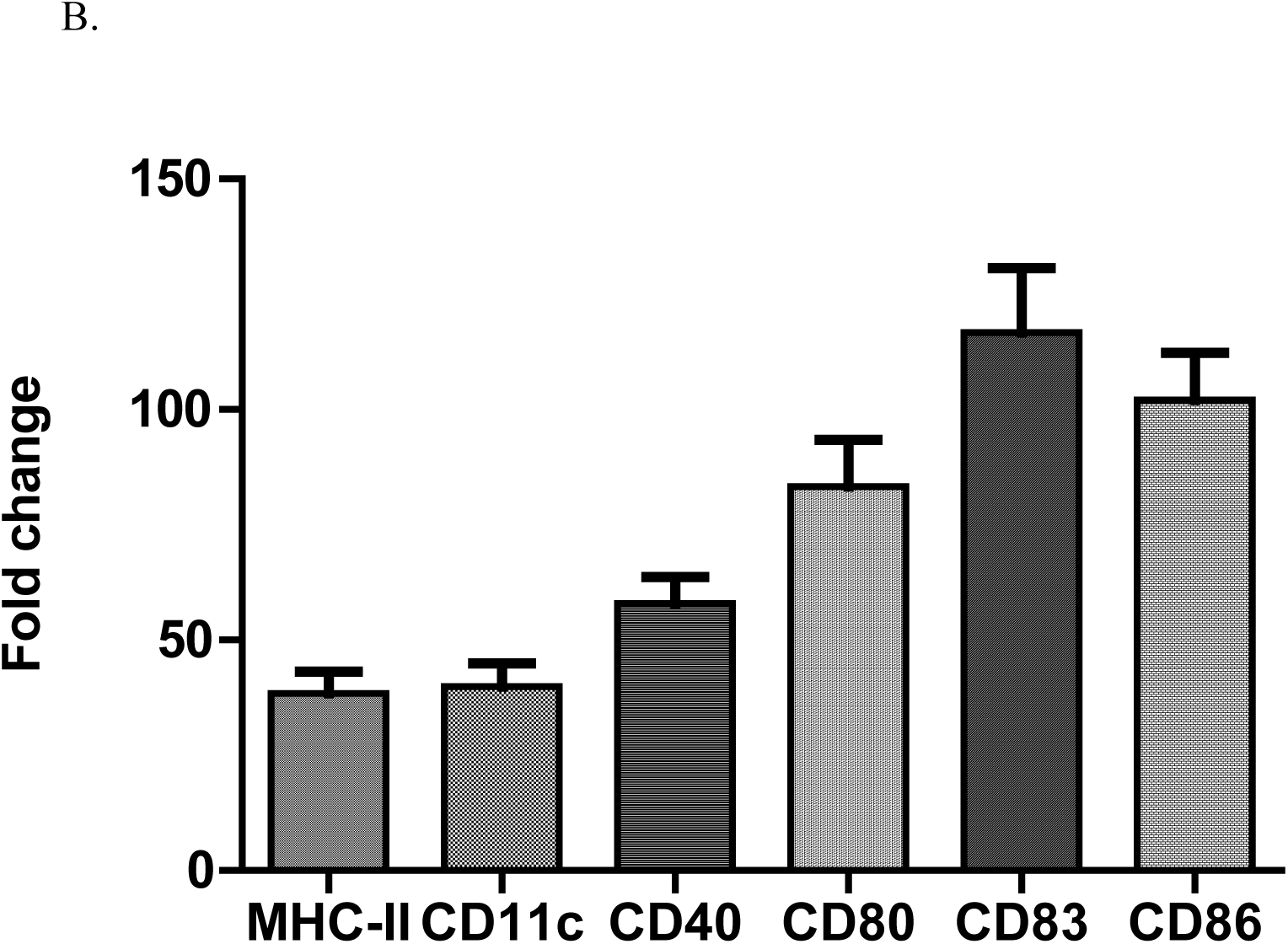
Comparative analysis of surface markers on immature and mature ckBM-DCs. Cells were cultured in the presence of 50 ng/ml GM-CSF + 50 ng/ml IL-4 for 6 days, and then stimulated with 500 ng/ml LPS for 30 hours. (A) Immature cells are on the left (A1,B1,C1) and mature cells are on the right (A2,B2,C2). Immunofluorescence analysis was performed using a FITC labeled mouse-anti-chicken MHC-II antibody (A1, A2). Cells were also stained with mouse anti-chicken CD11c (B1,B2) and mouse anti-chicken CD40 (C1,C2) followed by a goat-anti-mouse secondary. A representative image is shown for each at 100x magnification. (B) Cellular RNA was extracted to measure expression levels of surface markers in mature ckBM-DCs. RNA was normalized using the Ck 28S house-keeping gene. The data is expressed as the fold change in mRNA levels between immature (negative control) and mature ckBM-DCs for MHC-II, CD11c, CD40, CD80, CD83, and CD86. The data shown is a representative of three independent experiments. Error bars represent the standard deviation.

### 3.4 Immature ckBM-DCs retain the capability to phagocytosize foreign antigens

To test phagocytosis, 0.5-um carboxylate modified fluorescent red latex beads and FITC labeled-inactivated H5N9 avian influenza virus particles were added to immature ckBM-DCs. The cells were able to phagocytose both the beads (Fig. 4. A) and viral particles (Fig. 4. B). The beads and virus were observed in the cytoplasm (Fig. 4. A,B).

**Fig. 4.**
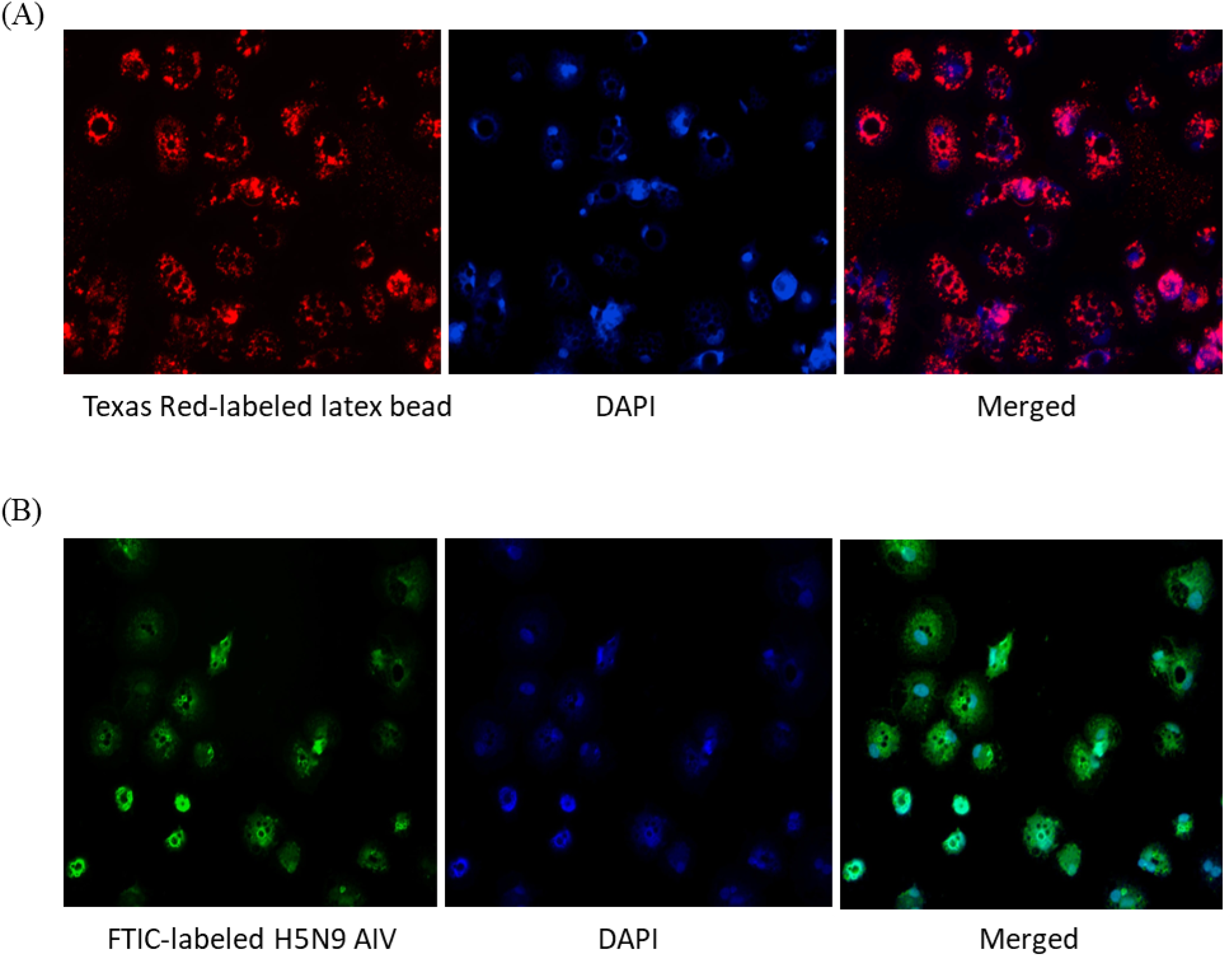
Functionality of immature ckBM-DCs. Cells were cultured in the presence of 50 ng/ml GM-CSF + 50 ng/ml IL-4 for 6 days. (A) ckBM-DCs were incubated with 0.5-um carboxylate modified fluorescent red latex beads or (B) FITC labeled-inactivated H5N9 avian influenza virus for 4 hours. Following incubation cells were counterstained with DAPI, washed 5x with PBS, and visualized with immunofluorescence microscopy. A representative image is shown for each at 100x magnification.

### 3.5 AIVs are capable of infecting ckBM-DCs

To test whether ckBM-DCs can successfully be infected with AIV, expression levels of SA-α2,3-Gal and SA-α2,6-Gal receptors on the cell surface were tested by immunofluorescence microcopy. Results demonstrated that both SA-α2,3-Gal (Fig. 5. A2) and SA-α2,6-Gal (Fig. 5. B2) receptors were extensively expressed in the cells, however the SA-α2,3-Gal produced higher levels of expression. Cytopathic effect (CPE) was observed at 20 hpi after infecting the cells with pandemic H1N1 (SA-α2,6-Gal preference) and H5N9 (SA-α2,3-Gal preference) (Fig. 5. C1, C2). Immunofluorescence microscopy with NP monoclonal antibodies and FITC linked α-mouse antibodies demonstrated the viruses were able to localize and replicate in the cells (Fig. 5. D1, D2)

**Fig. 5.**
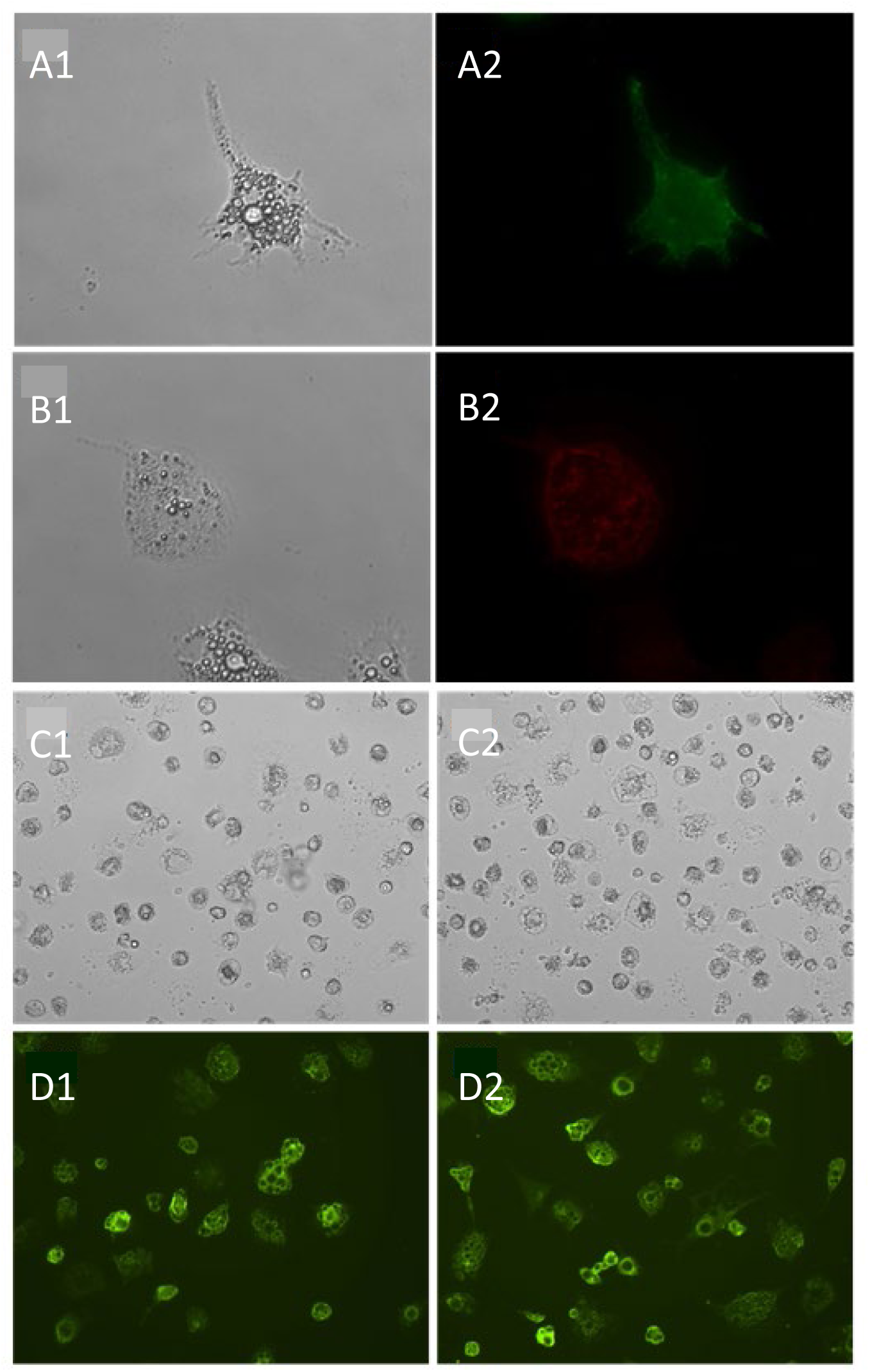
Distribution of sialic acid receptors on ckBM-DCs and susceptibility to pandemic H1N1 and H5N9 viruses. Immature ckBM-DCs (A1, B1) were stained with FITC-labeled MAA (SA-α2,3-Gal) (A2) or TRITC-labeled SNA (SA-α2,6-Gal) (B2), counter stained with DAPI, and visualized by immunofluorescence microscopy. CkBM-DCs were infected at an MOI of 1 with A/turkey/Virginia/SEP-4/2009 H1N1 (SA-α2,6-Gal preference) and A/turkey/Wisconsin/68 H5N9 (SA-α2,3-Gal preference). At 20 HPI, viral-infected cells, H1N1 (C1) and H5N9 (C2), were washed 2x with PBS, fixed with methanol, and observed by microscopy. Viral NP proteins, H1N1 (D1) and H5N9 (D2), were detected using a mouse-anti-NP antibody followed by a FITC-conjugated anti-mouse IgG secondary (D1,D2). A representative image is shown for each at 100x (A1,A2,B1,B2) and 200x (C1,C2,D1,D2) magnification.

### 3.6 The ckBM-DCs can be infected with both LPAIVs and HPAIVs

ckBM-DCs were infected with HPAIV and LPAIV of H5N2 and H7N3 subtypes to determine the effect on cell morphology and viral replication. At 8 hpi, there was little difference among the negative control and infected groups. At 8 hpi, all infected cells underwent some degree of morphological change, with H7N2 strains being more abundant in numbers of rounded cells compared to H5N2 strains. At 24 hpi, CPE was observed in the form of detachment of cells and change in morphology (rounding) of all infected cells, regardless of subtype or pathogenicity (Fig. 6. A). There was little difference in severity of CPE between the H5N2 strains, but cells infected with HPAI H7N3 demonstrated more severe levels of CPE with larger number of detached cells compared to LPAI H7N3. In terms of viral growth, LPAI H5N2 demonstrated a titer of 10^5.5^ EID_50_/ml at 24 hpi, compared to HPAI H5N3 which demonstrated a titer of 10^4.8^ EID_50_/ml. In contrast, HPAI H7N3 demonstrated a higher titer compared to LPAI H7N3, with titers of 10^6.5^ EID_50_/ml and 10^4.8^ EID_50_/ml, respectively (Fig. 6. B).

**Fig. 6.**
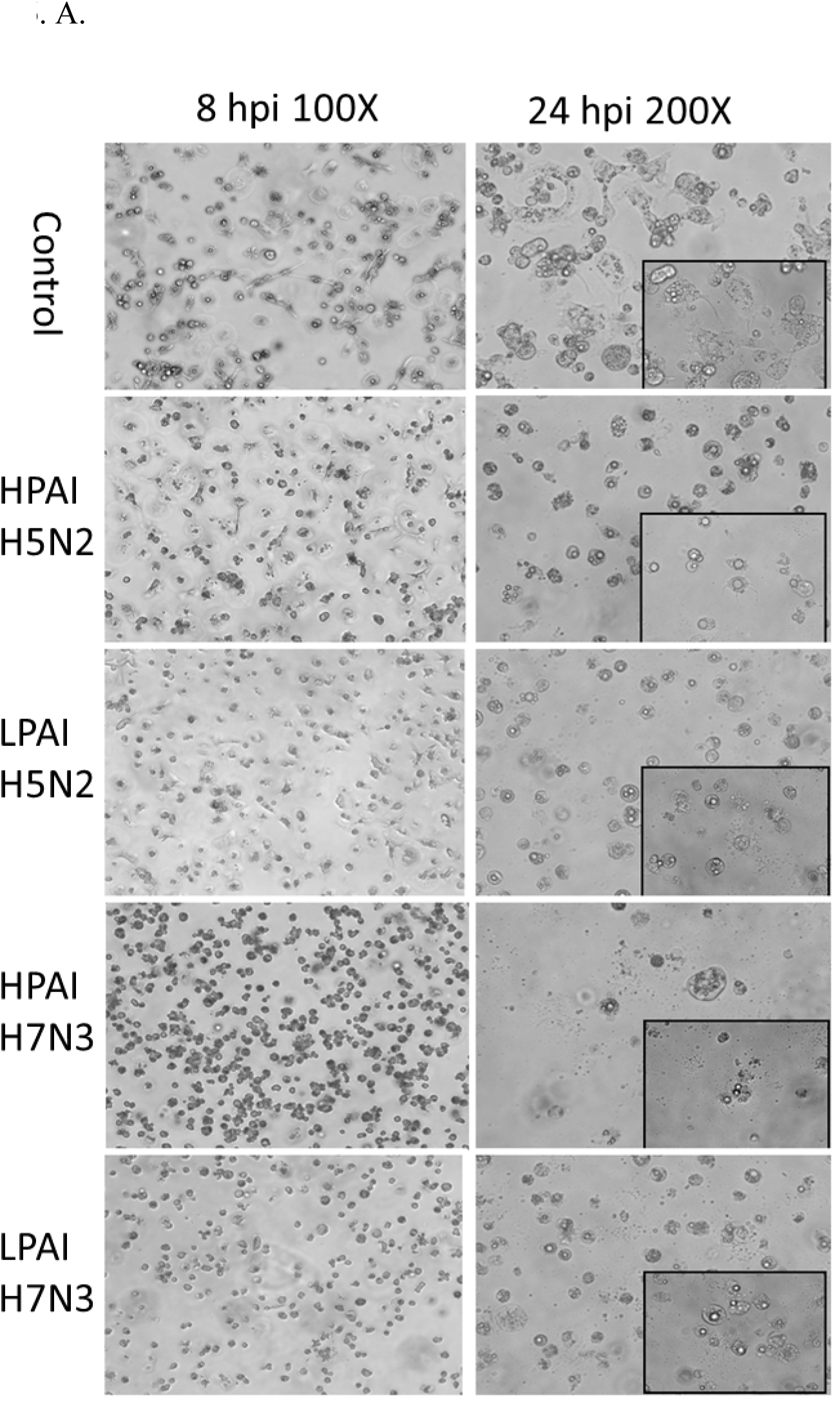
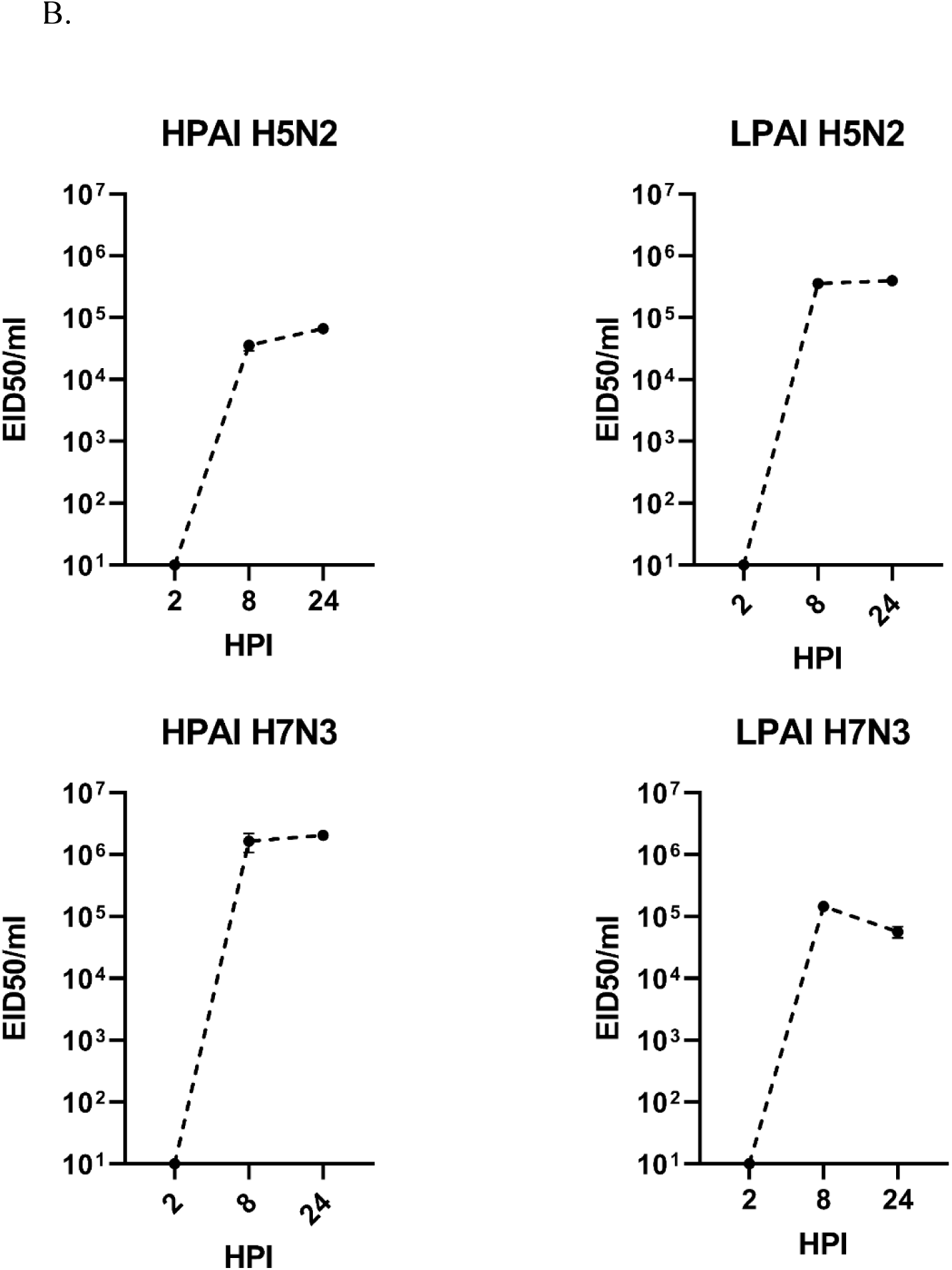
Change of morphology and growth of ckBM-DCs infected with LPAIV and HPAIV. ckBM-DCs were infected at a MOI of 1 with LPAIV (A/Chicken/Pennsylvania/21525/1983 H5N2, A/Cinnamon Teal/Mexico/2817/2006 H7N3) and HPAIV (A/Chicken/Pennsylvania/1370/1983 H5N2, A/Chicken/Jalisco/CPA1/2017 H7N3) viruses. (A) CPE and cellular morphological changes were observed by microscopy at 8 and 24 HPI. A representative image is shown for each at 100x and 200x magnification. (B) Supernatants were obtained at 2, 8, and 24 HPI and viral titers were evaluated by EID50. The data shown is a representative of three independent experiments. Error bars represent the standard deviation of triplicate samples.

### 3.7 ckBM-DCs infected with HPAIVs demonstrates higher expression of immune markers compared to LPAIV

Increased expression of IFN-α and Mx genes, which are indicative of a viral infection, were increased in all groups (Fig. 7.). All HPAI infected cells expressed significantly higher levels of both IFN-α and Mx genes, compared to their LPAI counterparts, between 20-500-fold, depending on virus and gene (Fig. 7. A). Increased expression of TLR receptors and MHC-I were observed in both HPAI and LPAI groups, with HPAI groups demonstrating higher expression levels compared to LPAI groups with a 10-700-fold difference. (Fig. 7. B). Similar trends were observed in gene expression levels of proinflammatory cytokines IL-1β, IL-6 and apoptotic genes Casp-3 and Casp-8, in which all HPAIV infected cells demonstrating significantly higher levels of expression compared to LPAI groups, between 20-800-fold difference. (Fig. 7. C, D).

**Fig. 7.**
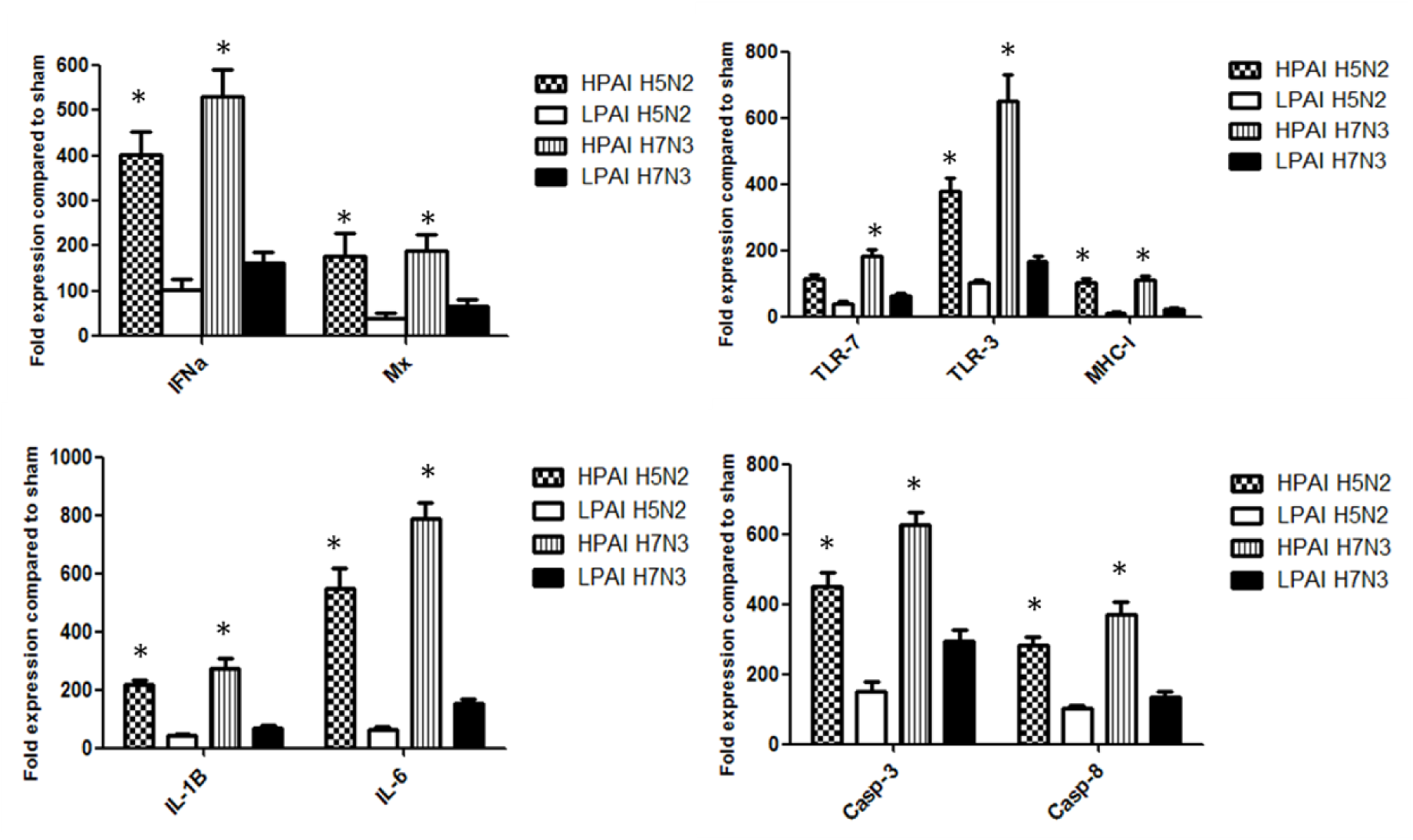
Cytokine expression levels of ckBM-DCs infected with HPAIV or LPAIV. ckBM-DCs were infected at a MOI of 1 with LPAIV (A/Chicken/Pennsylvania/21525/1983 H5N2, A/Cinnamon Teal/Mexico/2817/2006 H7N3) and HPAIV (A/Chicken/Pennsylvania/1370/1983 H5N2, A/Chicken/Jalisco/CPA1/2017 H7N3) viruses. Cellular RNA was extracted 8 HPI to measure relative gene expression levels of IFNα, Mx, TLR-7, TLR-3, MHC-1, IL-1B, IL-6, Casp-3, and Casp-8. RNA was normalized using the Ck 28S house-keeping gene. The fold change in expression was determined by the comparison of infected ckBM-DCs to sham-infected ckBM-DCs. Tukey one-way ANOVA analysis was performed to determine significant differences between LPAIV and HPAIV infected ckBM-DCs. * indicates a significant difference (p<0.05). The data shown is a representative of three independent experiments. Error bars represent the standard deviation of triplicate samples.

## 4. Discussion

The innate immune system plays a central role in detecting viral pathogens and mounting an early response by activating inflammatory and antiviral defense mechanisms. DCs are essential in controlling and bridging the gap between innate and adaptive immune responses due to their ability to process and present antigens, which is crucial for combatting virus infection. However, it is still largely unknown on how infection by AIVs can affect DCs, such as how morphology is changed or their immune profile is altered. This study has demonstrated that DCs were able to be infected by AIV, support viral growth, and phagocytosize viral particles even at the immature stage, which may promote antigen presentation. As proven in our study, DCs can be infected with both HPAIV and LPAIV. However, differences did exist between them in terms of morphological change and viral growth by subtype. While morphological changes were observed in all infected groups, there was little difference in severity of CPE between the H5N2 viruses at 8 and 24 hpi, however CPE was severe at 24 hpi for both viruses. In contrast, HPAI H7N3 demonstrated more severe CPE compared to the LPAI H7N3, demonstrating a correlation with pathogenicity and CPE. Viral titers also correlated with CPE and pathogenicity, with the titer of HPAI H7N3 demonstrating a 1.7 log difference in EID_50_/ml titers compared to LPAI H7N3. The results are consistent with a previous study in which HPAI H7N1 had higher levels of viral RNA compared to LPAI [25]. Interestingly, LPAI H5N2 replicated better in ckBM-DCs than HPAI H5N2, with a 0.7 log difference in terms of EID_50_/ml titers. While the exact reason is not clear, it may indicate a delayed replication or be attributed to a difference in subtypes. One study did report that a LPAI H7N1 replicated better in chicken lungs, suggesting the relationship of virus replication and pathogenicity may not always correlate [37].

Cell death is normally induced by apoptotic genes, usually by up-regulation of related caspase genes during viral infection [38] . This was demonstrated in our study, with Casp-3 and Casp-8 expression increasing in all infected groups, regardless of HPAI and LPAI. However, HPAI infected groups demonstrated significantly higher expression of Casp-3 and Casp-8 compared to their LPAI counterparts, indicating a correlation between caspase gene expression and pathogenicity. Studies has demonstrated AIV generally causes caspase-dependent apoptosis based on Casp-3 activation, which results in nuclear export of newly synthesized viral nucleoprotein (NP) and elevated virus replication, this suggests Casp-3 activation during onset of apoptosis is a crucial event for AIV propagation [39, 40]. One study has reported primary duck cells infected with LPAI H2N3 and classical H5N1 strains underwent rapid cell death compared to primary chicken cells, with both cells showing similar levels of viral RNA but less amount of infectious virus in duck cells [41]. Such rapid cell death was not observed in the same study with duck cells infected with a contemporary Eurasian H5N1 strain fatal to ducks, indicating the rapid apoptosis may be part of a mechanism of host resistance against AIV [42]. An increased expression of caspase genes demonstrated in our study may further support the notion that AIV can induce cell death via Casp-3 and Casp-8.

During AIV infection, ssRNA and dsRNA are recognized by a specific group of PRRs. In this study, HPAIV infected cells demonstrated significantly higher expression levels of TLR-3 and TLR-7 compared to cells infected with LPAIV. However, the level of TLR expression did not correspond to the amount of viral load as the titers had mixed results between the HPAI and LPAI strains. Furthermore, the TLR-3 expression levels were much higher in HPAI H7N3 compared to HPAI H5N2s. One study reported that TLR-3 expression levels significantly increased at 4 hpi and 16 hpi with HPAI H7N1 infections, whereas the level of increase in HPAI H5N2s were more gradual [25]. Another study reported STAT-3 expression was not adversely affected by LPAIV H3N2 in chicken cells, but expression levels were halved in chicken cells infected by HPAI H5N1 [42]. In contrast, STAT-3 expression levels were significantly elevated in duck cells, indicating infection with the same H5N1 strains had a less adverse effect. Thus, it can be speculated that differences in the cell signaling process may affect cytokine responses. Our results demonstrated that like the TLR gene expression profiles, expression levels of proinflammatory related genes were higher in the HPAI groups in the early stages of infection, compared to LPAI groups. Vervelde et al (2013) reported that levels of IFN-α were elevated in HPAI infected DCs and maintained between the early stages of infection up to 24 hpi, compared to the LPAI infected DCs where most of the IFN-α expression occurred in the early stages. Also, our study has demonstrated that the expression levels of IFN-α and IL-6 genes in DCs were higher in the HPAI H7N3 group, compared to the HPAI H5N2 group, suggesting the ability to activate host innate response may vary depending on the viruses and the host cells. Several studies have reported high levels of IL-6, IL-12 and IL-18 cytokine expression in the lungs and spleens of chickens infected with H5 HPAIVs while type 1 interferons were mostly present in the plasma and tissues [43–46]. Also, another study reported similar amounts of viral RNA and cytokine expression was detected in H7N1 in chickens, regardless of high and low pathogenicity [37]. Kuribayashi et al (2013), demonstrated that H7N1 strains can replicate more efficiently in chickens compared to H7N7, especially in the brain and are able to trigger excessive expression of inflammatory and antiviral cytokines, such as IFN-γ, IL-1β, IL-6, and IFN-α, in proportion to its proliferation. In contrast, another study reported that human-origin DCs infected with HPAI H7 resulted in delayed and decreased expression of cytokines, including type 1 interferons, compared to other AIV subtypes [47]. Thus, the difference of immune profiles of the host cell might be attributed to the specificity of the AIV. Furthermore, HPAI viruses may impair the regulatory activity of the TLR pathway, which is responsible for controlling the magnitude and duration of the inflammatory response, therefore leading to an uncontrolled immune response and cytokine storm. The acute uncontrolled innate immune response leading to overexpression of proinflammatory cytokines may be one of the causes for swift death in mammals infected with HPAI. Thus, one can speculate deregulation of these cytokines in chicken DCs may lead to multiple organ failure, as frequently seen in mammals.

Mx is a well-known antiviral protein, which can be induced by type 1 interferons [15]. However, susceptibility to the inhibitory effects of Mx may vary by strain and host [14, 48–50]. In this study, higher type 1 interferon expression levels were observed along with elevated expression of the Mx gene. However, despite the presence of elevated type 1 interferon and Mx expression levels, viral replication in DCs were not significantly inhibited. Also, presence of rapid cell death and activation of caspase-dependent apoptosis did not appear hinder the output of viral load. To date, the full complement of genes and their exact roles which contribute to antiviral properties are not well defined. However, one might speculate that the PRR dependent immune response may play a crucial role in mounting an antiviral defense, given the role of the TLR-7 and RIG-1 receptor signaling. For instance, it was shown that the presence of RIG-1 in cells stimulates expression of several key genes involved in innate immune response, which is crucial against viral infections such as influenza [51].

Overall, we were able to demonstrate that AIV can infect and replicate in chicken DCs regardless of pathogenicity. HPAI subtypes trigger a significantly higher expression of various immune factors compared to LPAI subtypes, suggesting a dysregulation of the immune system. The increase in DC activation following infection may be indicative of dysregulation of the immune response typically seen with high pathogenic avian influenza infections.

## Conflict of Interest

The authors declare that the research was conducted in the absence of any commercial or financial relationships that could be construed as a potential conflict of interest.

## Data availability statement

The raw data supporting the conclusions of this article will be made available by the authors, without undue reservation.

## Author contributions

**JM**: methodology, formal analysis, writing and revision. **KS** and **KC**: methodology, writing and revision. **KB** and **DK**: conceptualization, methodology, formal analysis, writing—original draft and preparation, and writing—review and editing. All authors contributed to the article and approved the submitted version.

## Funding

This work was funded by USDA-NIFA AFRI grant # 2015-67015-22968 as part of the joint USDA-NSF-NIH-UKRI-BSF-NSFC Ecology and Evolution of Infectious Diseases (EEID) program and USDA-ARS CRIS # 6040-32000-081-00D. This research was also supported in part by an appointment to the ARS Research Participation Program administered by the Oak Ridge Institute for Science and Education (ORISE) through an interagency agreement between the U.S. Department of Energy (DOE) and USDA, and by EEID. ORISE is managed by ORAU under DOE contract number DE-SC0014664. The findings and conclusions in this publication are those of the authors and do not be necessarily represent the official policy of the USDA, DOE, or ORAU/ORISE. Any use of trade, product or firm names is for descriptive purposes and does not imply endorsement by the U.S. Government. The USDA is an equal opportunity provider and employer.

## Acknowledgments

The authors thank Dr. Hai Jun Jiang and Ryan Sweeney for excellent technical assistance.

## Notes

### Competing Interest Statement

The authors have declared no competing interest.

